# Effect of white mulberry (*Morus albas* L.) On common carp performance, biological parameters, and blood picture

**DOI:** 10.1101/2021.06.01.446541

**Authors:** Nasreen Mohi Alddin Abdulrahman

## Abstract

This study has been done for evaluating the growth performance and health performance of *Cyprinus carpio* isoenergetic with isonitrogenous diets which contain three levels (0, 10, 15, gm of mulberry fruit powder, have been used for a period of 12 weeks. The effect of white mulberry (*Morus albas* L.) fruit powder on common carp, T2 was significantly higher than other treatments in each of growth performance and feed utilization parameters. the adding of white mulberry fruit powder significantly increased each of CBC counts in T2 of RBC and HCT%. The conrol was significantly higher in each of MCH, MCHC, MCV and Platelets count. Cholestrol, triglyceride and HDL were higher significantly in T2, LDL was significantly higher in control. The white mulberry (*M. albas* L.) fruit powder effect significantly in, GPT, GOT, in T2 and the CKI and blood suger were higher significantly in the control, the control was significantly higher in each of total protein, and albumin ratio. T2 was higher significantly in Intestine Length Index in both fish weigth and length, and both fish weigth without viscera and/ or head. No significant differences in each of Intestine Weight Index among treatments, blood Globulin ratio, Hemoglobin count and VLDL.

## Introduction

Mulberry (Morus spp.) is a tropical and subtropical fruit plant that belongs to the genus Morus, the tribe Moracea, and the family Moraceae (1). This species’ range includes East, West, and South East Asia, South Europe, South of North America, Northwest of South America, and parts of Africa. White mulberry, black mulberry, and red mulberry are the most widely grown plants in the *Morus* genus (2).

*Morus albas* L. (Moraceae) Parts of the white mulberry plant have been used as medicine in Chinese medicine for a long time. White mulberry is cultivated through leaves is the main food source for silk worms. Traditionally, the white mulberry plant has been used to treat weakness, fatigue, anemia, and preastare grayi in continence, tinniTs, dizziness and constipation in the alderly patient so far include analgesic, antiasthmatic, antirhaumatic, antiTssive, as expectorant, hypotensive and brain tonic (3).

Özgen et al. (2) stated that mulberry had essential effects in human health with the help of its antioxidant contents. The demand for fruit species containing anthocyanins and anthocyanidins has been increasing due to the identification of flavonoids having anticarcinogenic effects in the sTdies in recent years. Mulberry is also included in this fruit species. In the study of Gundogdu et al., (4) The chlorogenic acid and antioxidant contents of M. nigra fruit were found to be substantially higher than those of M. alba and M. rubra. It has been documented that phenolic compounds such as chlorogenic acid and catechins, which are commonly used, have antimutagenic and anticarcinogenic properties. Furthermore, cinnamic acids, a sub-group of nitrosamines suspected of causing cancer, are used to block them (5).

Mulberry is also used as a medicinal plant, with the biologically active pharmacokinetic compounds present in the leaf, stem, and root sections being used to strengthen and enhance human life. Industrialists have been involved in more industrial processing of mulberry through the preparation of various goods in the pharmaceutical, food, cosmetic, and health care industries. As mulberry is being exploited by sericulture, pharmaceutical, cosmetic, food and beverage industries along with its utilization in environmental safety approach; it is appropriate to call it as a most suitable plant for sustainable development (5). So the aim of using *M. alba* L to examine its ability to enhance common carp growth, performance, blood picture and meat quality.

## Materials and Methods

*M. alba* will be used with different levels were incorporated to a commercial diet to form three diets as follows: T1; a commercial diet that does not contain any supplements, T2 a commercial diet enriched with 10 g/Kg, T3 with 15 g/Kg diet.

Group: In fish diet 0% *Morus* fruit,

Group 2: In fish diet 10 g *Morus* fruit/ kg diet,

Group 3: In fish diet 15 g *Morus* fruit/ kg diet,

The chemical composition of the used diet is shown in Table 1.

**Table 1:** Chemical composition of control diet treatment.

| Ingredient | T1 ( Control ) |
| --- | --- |
| Crude Protein % | 29 |
| Crude Fat % | 2.2 |
| Dry Matter | 87.4 |
| Crude Fiber % | 4.7 |

### Common carp

*C. carpio* L. was transported from a local earthen pond in Daqoq, Iraq, and acclimated to laboratory conditions for three weeks, during which time the fish were fed a commercial diet (28 percent crud protein). Following that, 135 fish (95–105 g) were distributed at a density of 15 fish per tank in nine 100-L tanks. Air stones connected to an air pump supply compressed air to each tank. For 12 weeks, fish were fed one of the diets in triplicates up to apparent satiation three times a day at 9:00, 13:00, and 17:00 h. Every day, half of each tank’s water, along with the fish faces, will be drained and replaced with well-aerated tap water from a storage tank.

### Growth performance

Fish from each tank were captured, counted, and weighed at the end of the experiment. The following parameters of fish growth and feed use will be calculated:

Weight gain (g) = W2-W1

Specific growth rate (SGR; %g/day) = 100 [Ln W2 (g) -Ln W1 (g)]/T;

where W1 is initial weight (g), W2 is final weight (g), and T is the feeding period (day); Relative growth rate was calculated according to the method described by Brown, (1957). Relative growth rate (RGR %) = Weight gain/Initial weight x 100

= W2–W_1_/ W_1_ x 100

Specific growth rate was calculated according to the method described by Uten, (1978). Specific growth rate (SGR) % = (Ln final body weight–Ln initial body weight)/ experimental period) x 100

= ((Ln W2–Ln W1)/ T) x 100

Feed conversion ratio (FCR) = Total feed fed (gm.)/ Total wet weight gain (g) according to Feed efficiency ratio was calculated as previously described by Uten, (1978)

FCR= Total feed/ Total wet weight gain

Feed efficiency ratio (FER) = Total weight gain (gm.)/ Total feed fed (gm.)

Protein efficiency ratio was calculated as:

Protein efficiency ratio (PER) = Total wet weight gain (gm/fish)/ Amount of protein fed (gm./fish).

Fat efficiency ratio (FER) = Total wet weight gain (gm/fish)/ Amount of fat fed (gm./fish).

### Biological parameters

At the end of the experimental period, five fish were randomly taken from each tank, anaesthetized with clove powder (30 mg/L) and blood samples will be collected from the caudal vein. After that, fish weight and length measured; fish then dissected and liver, gills, viscera, and kidney will removed and weighed.

The organs-somatic indices of fish were calculated as follows:

Condition (K) factor = 100 (fish weight; g)/(fish length; cm)3 ;

Hepatic somatic index (HSI, %) = 100 (liver weight (g)/fish weight (g));

Gills somatic index (GSI, %) = 100 (gills weight (g)/fish weight (g));

Kidney somatic index (KSI, %) = 100 (kidney weight (g)/fish weight (g));

Spleen somatic index (SSI, %) = 100 (kidney weight (g)/fish weight (g)).

Intestine Length Index (According to fish Length) = 100 (Intestine length (cm)/fish length (g)).

Intestine Length Index (According to fish Weight) = 100 (Intestine length (cm)/fish weight (g)).

Intestine Weight Index = 100 (Intestine length (cm)/fish weight (g)).

Weigth without Viscera Index = 100 (Fish weight / Fish Weigth without Viscera Wt. without Viscera & Head = 100 (Fish weight / Fish Weigth without Viscera & Head

### Blood parameters

Blood samples from the caudal vein were split into two classes of Eppendorf Tbes. One set contained sodium heparinate (20 U/L), as an anticoagulant, for measuring white blood cells (WBCs), red blood cells (RBCs), hemoglobin (Hb), and hematocrit (Hct) (CBC counts parameters). The second group was given no anticoagulant and permitted to clot at 4°C before being centrifuged at 5000 g for 20 minutes at room temperature to obtain serum for testing the various biochemical parameters. Seruum biochemical analyses including glucose, total protein, total cholesterol, high density lipoprotein cholesterol (HDL), low density lipoprotein cholesterol (LDL), triglycerides, aspartate aminotransferase (AST), and alanine aminotransferase (ALT) were determined using an automatic biochemical analyzer using commercial kits from Spinreact, S.A. (Gerona, Spain).

Prior to statistical analysis, all data tested for normality of distribution using the Kolmogorov–Smirnov test. The homogeneity of variances among different treatments will be tested using Bartlett’s test. Then, data will subjected to one-way ANOVA to evaluate effects of different feed supplementation. Duncan’s test is used as a post hoc test to compare between treatments at P < .05. All the statistical analyses done via SPSS program version 20 (SPSS, Richmond, VA, USA).

## Results

Table 1 show the effect of white mulberry (*Morus albas* L.) fruit powder on common carp, T2 with 15 g/kg was significantly higher than other treatments in each of growth performance and feed utilization parameters.

**Table 1:**
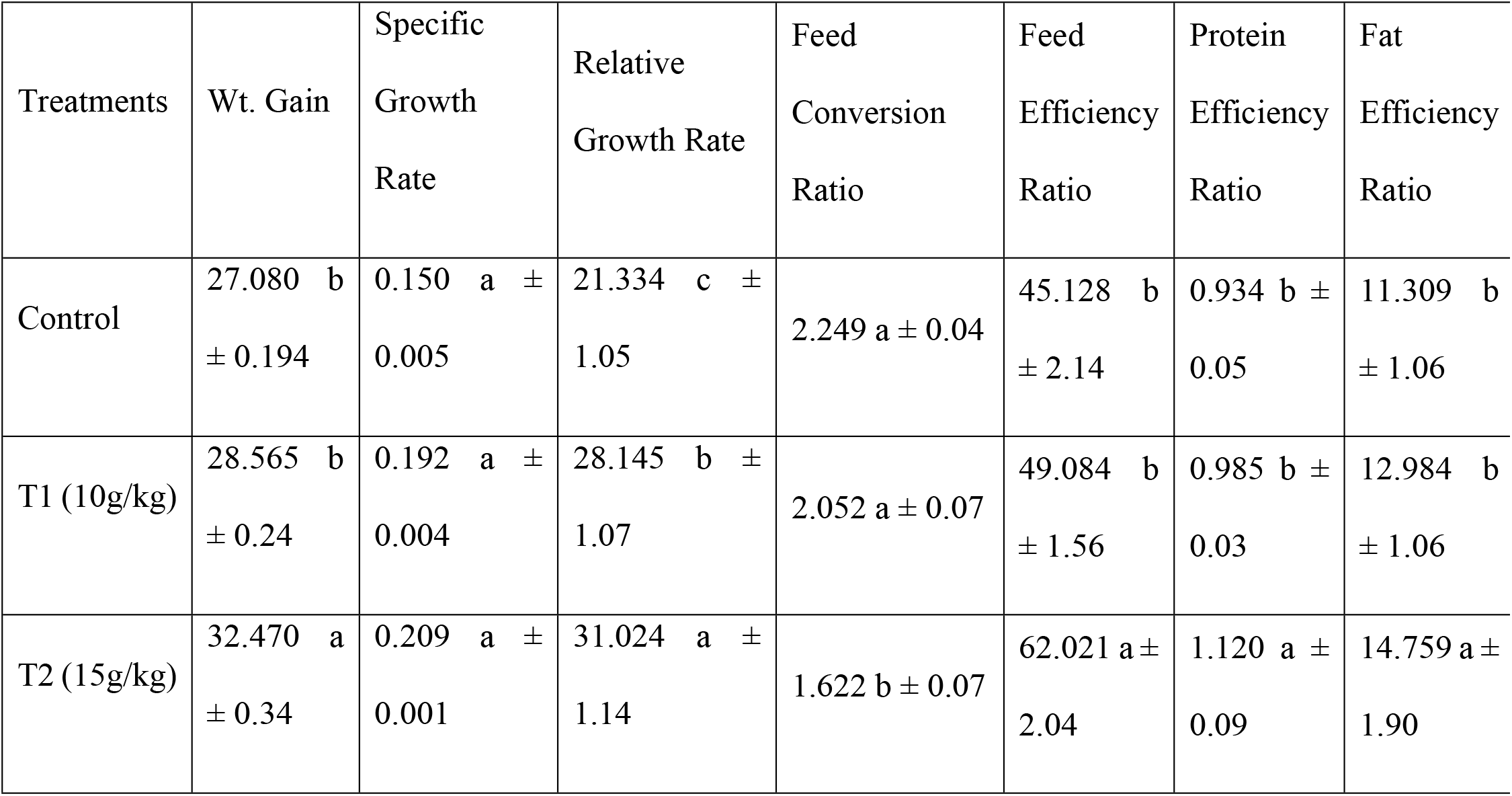
Effect of white mulberry (Morus albas L.) fruit powder on common carp growth performance and feed utilization

Different letter in same rows mean significant differences (p < 0.05)

Table 2 show that the adding of white mulberry (*M. albas* L.) fruit powder significantly increased each of CBC counts in T2 with 15g/kg diet of RBC and HCT(PCV)%. The conrol was significantly higher in each of MCH, MCHC, MCV and Platelets count. No significant differences observed in Hemoglobin count.

**Table 2:**
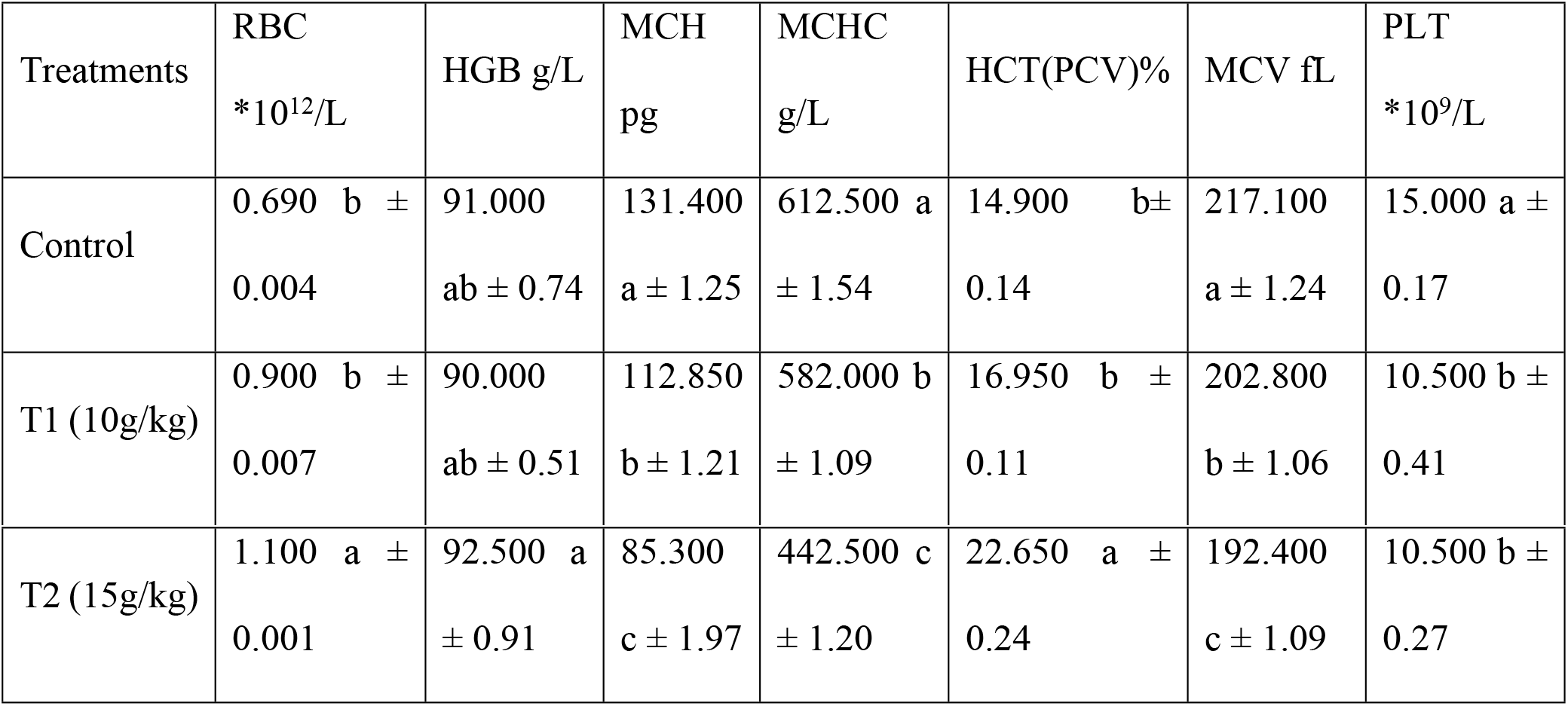
Effect of white mulberry (Morus albas L.) fruit powder on common carp blood picture

Different letter in same rows mean significant differences (p < 0.05)

Cholestrol, triglyceride and HDL were higher significantly in T2, LDL was significantly higher in control, no significant dfferences observed in VLDL as seen in table 3.

**Table 3:** Effect of white mulberry (Morus albas L.) fruit powder on common carp blood lipid profile

| Treatments | Cholesterol | triglyceride | LDL | HDL | VLDL |
| --- | --- | --- | --- | --- | --- |
| Control | 88.250 b ±0.15 | 181.000 bc<br>±0.14 | 20.750 a<br>±1.11 | 38.700 c<br>±0.04 | 38.855 a<br>±0.01 |
| T1<br>(10g/kg) | 89.000 b ±0.14 | 185.400 b<br>±0.12 | 17.600 b<br>±0.97 | 46.450 b<br>±0.01 | 37.100 a<br>±0.01 |
| T2<br>(15g/kg) | 93.650 a ±0.11 | 194.250 a<br>±0.12 | 17.800 b<br>±0.92 | 56.150 a<br>±0.01 | 36.200 a<br>±0.01 |

Different letter in same rows mean significant differences (p < 0.05)

The white mulberry (*M. albas* L.) fruit powder effect significantly in, GPT, GOT, in T2 and the CKI and blood suger were higher significantly in the control, the control was significantly higher in each of total protein, and albumin ratio, no significant in Globulin ratio as observed in table 4.

**Table 4:**
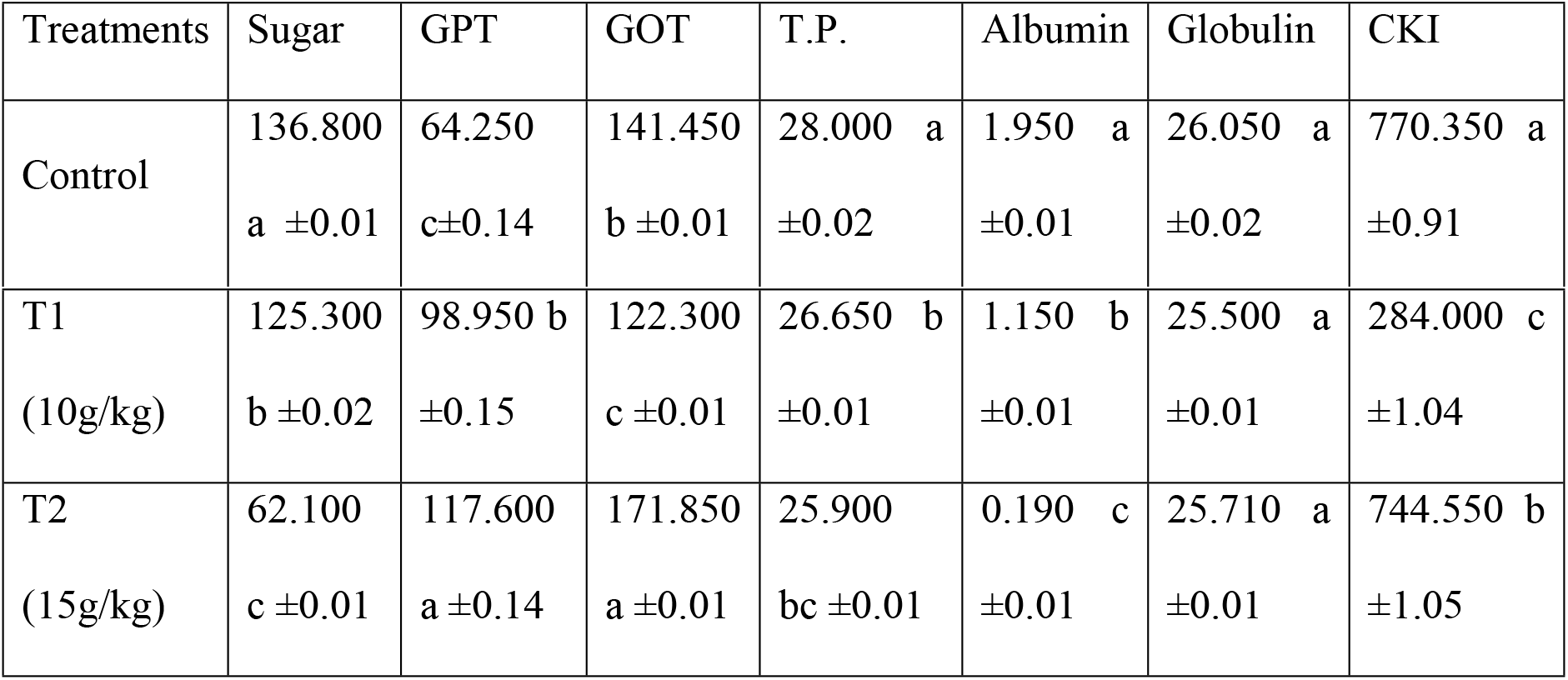
Effect of white mulberry (Morus albas L.) fruit powder on common carp blood biochemicals

| Treatments | Sugar | GPT | GOT | T.P. | Albumin | Globulin | CKI |
| --- | --- | --- | --- | --- | --- | --- | --- |
| Control | 136.800<br>a ±0.01 | 64.250<br>c±0.14 | 141.450<br>b ±0.01 | 28.000 a<br>±0.02 | 1.950 a<br>±0.01 | 26.050 a<br>±0.02 | 770.350 a<br>±0.91 |
| T1<br>(10g/kg) | 125.300<br>b ±0.02 | 98.950 b<br>±0.15 | 122.300<br>c ±0.01 | 26.650 b<br>±0.01 | 1.150 b<br>±0.01 | 25.500 a<br>±0.01 | 284.000 c<br>±1.04 |
| T2<br>(15g/kg) | 62.100<br>c ±0.01 | 117.600<br>a ±0.14 | 171.850<br>a ±0.01 | 25.900<br>bc ±0.01 | 0.190 c<br>±0.01 | 25.710 a<br>±0.01 | 744.550 b<br>±1.05 |

Different letter in same rows mean significant differences (p < 0.05)

Table 5 show the effect of white mulberry in some physiobiological parametrs without any significancy.

**Table 5:** Effect of white mulberry (*Morus albas* L.) fruit powder on common carp biophysiological indices

| Treatments | Hepatosomatic index | Spleen somatic index | Gill somatic index | Kidney somatic index | Condition Factor |
| --- | --- | --- | --- | --- | --- |
| Control | 1.476 a ±0.02 | 0.163 a<br>±0.04 | 6.104 a<br>±0.01 | 0.307 a<br>±0.01 | 1.528 a<br>±0.01 |
| T1 (10g/kg) | 1.672 a ±0.01 | 0.194 a<br>±0.05 | 5.205 a<br>±0.01 | 0.459 a<br>±0.01 | 1.412 a<br>±0.01 |
| T2 (15g/kg) | 1.807 a ±0.02 | 0.132 a<br>±0.04 | 6.282 a<br>±0.01 | 0.373 a<br>±0.01 | 1.610 a<br>±0.02 |

Different letter in same rows mean significant differences (p < 0.05)

No significant differences observed in Intestine Weight Index among treatments, T2 was higher significantly in Intestine Length Index in both fish weigth and length as seen in table 6.

**Table 6:** Effect of white mulberry (*Morus albas* L.) fruit powder on common carp intestine indices

| Treatments | Intestine Weight Index | Intestine Length Index (According to fish Weight) | Intestine Length Index (According to fish Length) |
| --- | --- | --- | --- |
| Control | 1.730 a ±0.02 | 19.696 b ±0.03 | 139.642 b ±1.24 |
| T1 (10g/kg) | 2.043 a ±0.02 | 22.359 ab ±0.03 | 139.524 b ±1.24 |
| T2 (15g/kg) | 2.339 a ±0.02 | 23.541 a ±0.02 | 158.182 a ±1.22 |

Different letter in same rows mean significant differences (p < 0.05)

Fish weigth in both without viscera and/ or head were higher significntly in T2 as shown in tablw 7.

**Table 7:**
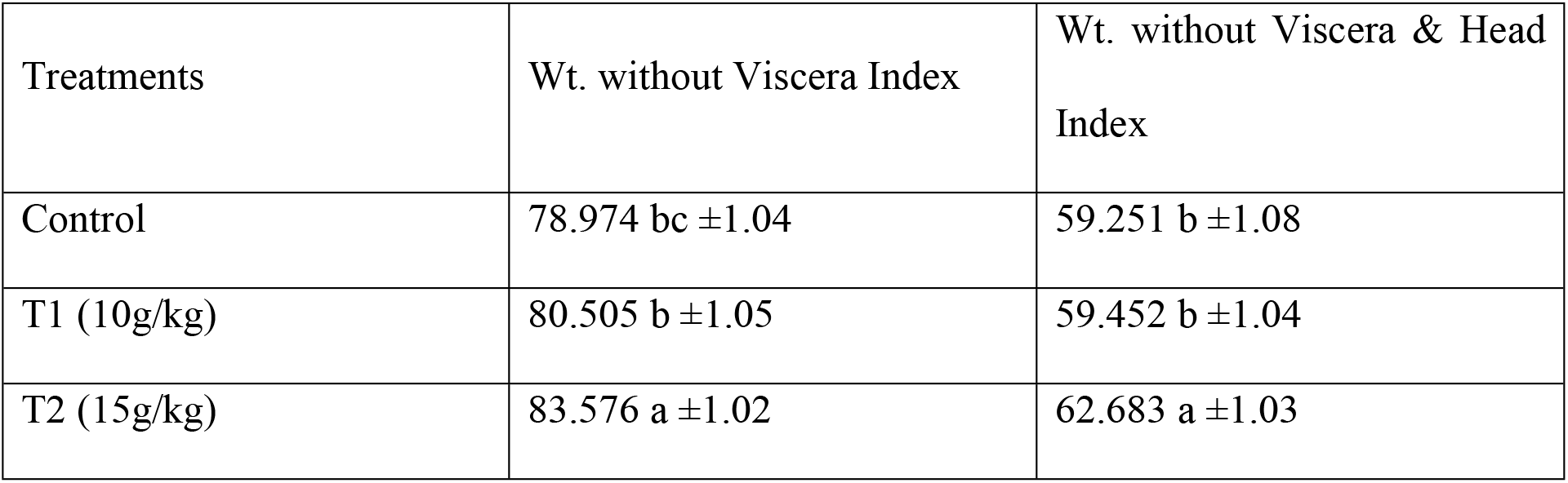
Effect of white mulberry (*Morus albas* L.) fruit powder on common carp meat indices

Different letter in same rows mean significant differences (p < 0.05)

## DISCUSSION

The current study was conducted for the first time to assess the effects of dietary supplementation with white mulberry fruit powder on common carp growth efficiency, blood parameters, and some biological parameters. Mulberry’s effectiveness has been attributed to flavonoids compounds, which can aid in improving fish health and performance (6). Despite the fact that mulberry is commonly used as a medicinal herb around the world, there have been no brief studies on its use as a feed additive in aquaculture.

The ability to acess the potential of using feed additives in aquaculture is based on growth success and feed conversion ratio. We discovered that dietary mulberry improved the weight gain, specific growth rate, and feed conversion ratio of common carp in the current research. Fish fed a diet supplemented with mulberry showed a substantial increase in weight gain while simultaneously decreasing the FCR. This result is consistent with previous studies that found that adding medicinal herbs to a diet improved growth-related parameters while decreasing the feed conversion ratio (7, - 11).

Shekarabi et al., (12) show that the SGR, WG, and SR values of rainbow trout fed diets supplemented with varying amounts of mulberry juice powder improved dramatically in a dose-dependent fashion following dietary supplementation with mulberry juice powder, with the highest values of fish fed 0.75 percent. When fish were fed mulberry juice powder, the feed conversion ratio (FCR) was also reduced. Li *et al.* (13) reported the dietary herbs enhance WG and FCR by enhancing nutrient utilization and enabling the intestinal flora’s functionality. The current study showed that integrating mulberry into the diet had a major effect on common carp growth and feed quality by growing FCR. Bioactive compounds such as anthocyanins, flavonols, and ellagitannins have been found in berry plants (14). Various active compounds in herbs promote digestion and improve protein assimilation in the intestinal tract of fish (15; 16; 14).

Blood biochemical and physiological indicators, such as serum compounds, may be used to detect possible beneficial effects of natural feed additives on aquatic animal welfare (17). In fish, blood cortisol and glucose levels are essential markers of environmental stress (18). Increased cortisol levels induce elevated blood glucose levels, which facilitate gluconeogenesis in the liver (19). When mulberry-fed fish were compared to the control group, glucose levels were found to be lower. Mulberry has an antistress effect, inhibiting the amplitude of elevated cortisol, thus reducing the degree of elevated glucose levels in blood, according to the slightly lower glucose levels in blood.

The findings also revealed that of the three amounts of dietary mulberry inclusion included in the analysis (0, 10, and 15 g/kg), the 15 g/kg dietary supplement was the most effective in enhancing efficiency. Several herbal plants also were tested for their growth promoting activity in aquatic animals such as Date Palm Seeds (Phoenix dactyliferous L.) by Ahmed et al., (20). Monitoring indices such as hematological parameters may show the status of the immune system in fish. However, in the current research, high doses of dietary mulberry (15 g/kg) increased the blood parameter to the amount shown in the control treatments. Mulberry fruit also has a range of biological properties, including antioxidant, neuroprotective, antiatherosclerosis, immunomodulatory, antitumor, antihyperglycemic, and hypolipidemic impact. Natural food ingredients can be nutritious and environmentally friendly with no harmful side effects. According to one study, polyphenol-rich mulberry can provide effective defense against EC-induced cytotoxicity and oxidative stress (22).

Mulberry extract has an important impact on the survival rate, stage development, and mortality rate of blue swimming crab larvae due to molting syndrome, according to Fujaya *et al.* (23). The higher the dose of ME in the artificial food, the higher the survival rate and the lower the mortality rate due to molting syndrome. These studies indicated that irrespective of the lutein and β-carotene content of the mulberry leaves in the diet, the fancy carp can convert them to astaxanthin in serum (24). The carotenoids in both of the diets were metabolized to astaxanthin, resulting in higher serum levels of this compound. This study proposes an innovative approach to replacing synthetic carotenoids in aquaculture diets by using native plant leaves (24).

---

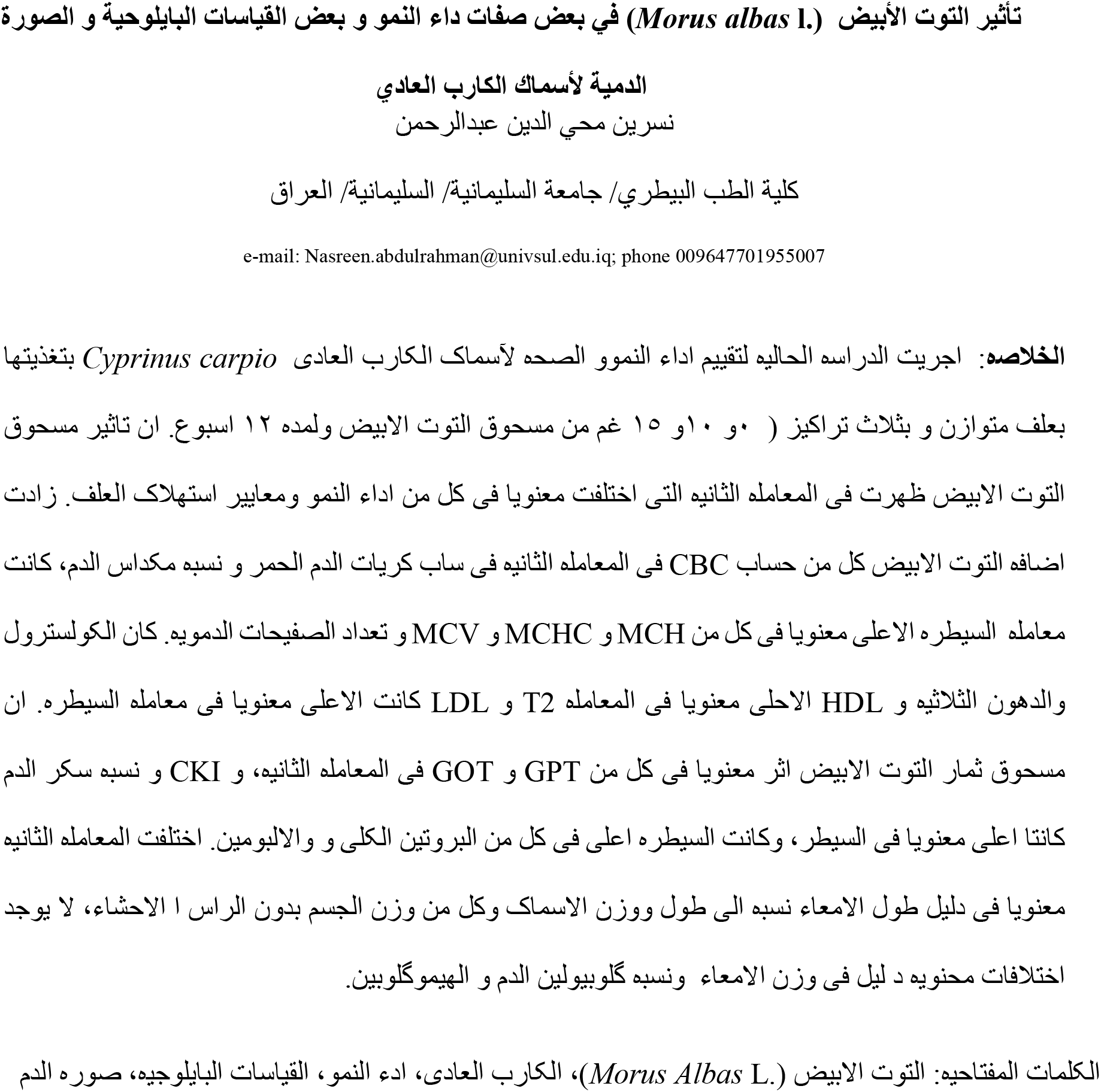

